# StratPal: An R package for creating stratigraphic paleobiology modeling pipelines

**DOI:** 10.1101/2024.11.22.624834

**Authors:** Niklas Hohmann, Emilia Jarochowska

**Affiliations:** Faculty of Geosciences, Department of Earth Sciences Utrecht University, The Netherlands

**Keywords:** Stratigraphy, Paleontology, Paleobiology, Stratigraphic Paleobiology, Mass extinction, Trait evolution

## Abstract

1. The fossil record is an important source of information to understand biological processes that take place over timescales not accessible to human observation or experiments. Fossil data is not a perfect reflection of past biological change, but a joint expression of stratigraphic, ecological, evolutionary, and taphonomic effects. Stratigraphic biology is a field dedicated to identifying these effects and accounting for them.
2. We present `StratPal`, a R package in which stratigraphic, ecological, evolutionary, and taphonomic modules can be combined into modeling pipelines for stratigraphic paleobiology.
3. We describe the types of data that can be modified and transformed using this approach, and briefly discuss potential extensions.
4. As working examples, we show how the pipeline can be used to model clustering of last occurrences at hiatuses and condensation surfaces, leading to artefactual “extinction events” caused by stratigraphic gaps.

## Introduction

The fossil record is key to understand biological change over timescales not accessible to human observation and experimentation. However, fossil data is not an unbiased representation of biological change, but a joint expression of stratigraphy, preservation, ecology, and evolution. Stratigraphic paleobiology is a new field which arose from the realization that information from all these disciplines needs to be integrated to recover biological information from the fossil record. Accounting for the structure of the geological record fundamentally improves reconstructions of trait evolution, phylogenetics, organism distribution and abundance, or origination or extinction dynamics from fossils (Holland and Patzkowsky 1999; 2015; Zimmt et al. 2021; Nawrot et al. 2018; Jarochowska et al. 2018; Danise et al. 2019; Hohmann 2021).

Stratigraphy investigates how time is represented in the physical rock record (strata). In empirical studies, disentangling the factors that contribute to the stratigraphic distribution of fossils is challenging, as a lot of data is required to constrain both biotic and abiotic parameters (e.g., age constraints, water depth, niches, and evolutionary change) (Dominici, Danise, and Benvenuti 2018; Warren Huntley and Scarponi 2015; Nawrot et al. 2018). In the fossil record, these parameters cannot be observed directly and need to be estimated, often with a large error. As a result, simulation studies are common: They provide “perfect knowlege” scenarios where all parameters are known. This allows to study the interaction of multiple effects *in silico*, thereby providing important stratigraphic null hypotheses against which empirical observations can be compared (Holland and Patzkowsky 2015; Zimmt et al. 2021).

A wide range of environments have been simulated in stratigraphic paleobiology, including sandy shelf (Holland and Patzkowsky 1999; Hannisdal 2006), carbonate (Hohmann, Koelewijn, et al. 2024), and terrestrial systems (Holland 2022). In these studies, biological change is cast against the backdrop of sedimentological and stratigraphic scenarios created through forward modeling. Each of these studies is idiosyncratic, making it difficult to identify general patterns or compare results due to methodological differences.

Here, we present the `StratPal` package, an R Software package that allows to combine taphonomic, ecological, evolutionary, and stratigraphic effects in a modular fashion to build modeling pipelines for stratigraphic paleobiology across environments (Hohmann 2024b; R Core Team 2024). The package provides utility functions to simulate trait evolution and event type data such as fossil ages or locations and first or last occurrences of taxa in the stratigraphic record, which can be used standalone or as part of a simulation pipeline. The package comes with extensive documentation in the form of vignettes (long form documentation and worked examples) that explain how modeling pipelines can be build, developer documentation, and example data of stratigraphic architectures simulated using CarboCAT (Burgess 2013, 2023).

As a worked example, we use the package to show how clusters of last occurrences and fossil accumulations form at predictable stratigraphic locations associated with gaps and stratigraphic condensation (i.e., a slowdown of sediment accumulation (Hohmann and Jarochowska 2023)), and how this effect changes along the onshore-offshore gradient in a carbonate platform. We find that, although clusters of last occurrences are generated by both gaps (hiatuses) in the record and by stratigraphic condensation, fossil accumulations appear only at intervals with stratigraphic condensation. This is because organismal remains formed during erosional intervals will generally be destroyed, while last occurrences are observed after destruction in the stratigraphic domain and thus cannot be removed by erosion.

The `StratPal` package provides a modular, unified approach to numeric stratigraphic paleobiology, allowing to build pipelines of varying complexity to study stratigraphic, taphonomic, ecological, and evolutionary effects in isolation or conjunction across depostional environments. Due to its modular nature, it can be linked with outputs from any sedimentary forward model (e.g., CarboCAT (Burgess 2013), CarboKitten (Hidding et al. 2024), or SedFlux (Hutton and Syvitski 2008)), thus facilitating comparisons across depositional environments. The package integrates with the existing package ecosystem for analytical paleobiology (e.g., `admtools` (Hohmann 2024a) or `paleoTS` (Hunt 2006; 2022), and provides extensive user and developer documentation to facilitate integration with other R packages. It is Open Source and encourages contributions by other researchers through appropriate licensing, contributing instructions, and development with version control.

## The StratPal package

### Conceptual framework

The `StratPal` package provides (1) functions to simulate trait evolution and fossil occurrence data, and (2) functions that act as filters on the original signal and modify it to emulate the effects of taphonomy, stratigraphy, and niche affinity (Figure 1). Importantly, the original biological signal is simulated in the time domain (*T* dimension, SI units s and derived units yr or Myr), but is observed in the stratigraphic domain (*L* dimension, SI units m) (Hohmann, De Vleeschouwer, et al. 2024). The stratigraphic domain is in length dimension following the convention in paleobiology, whereby strata in a geological section or core are treated as laterally uniform and perfectly sampled and change is only observed between strata but not within them. Under this convention, fossil observations can be treated as one-dimensional, even though in practice this simplification is often violated. The transformation between the time and stratigraphic domains is performed via age-depth models that associate points in time with stratigraphic positions and *vice versa* (Hohmann 2024a).

**Figure 1:**
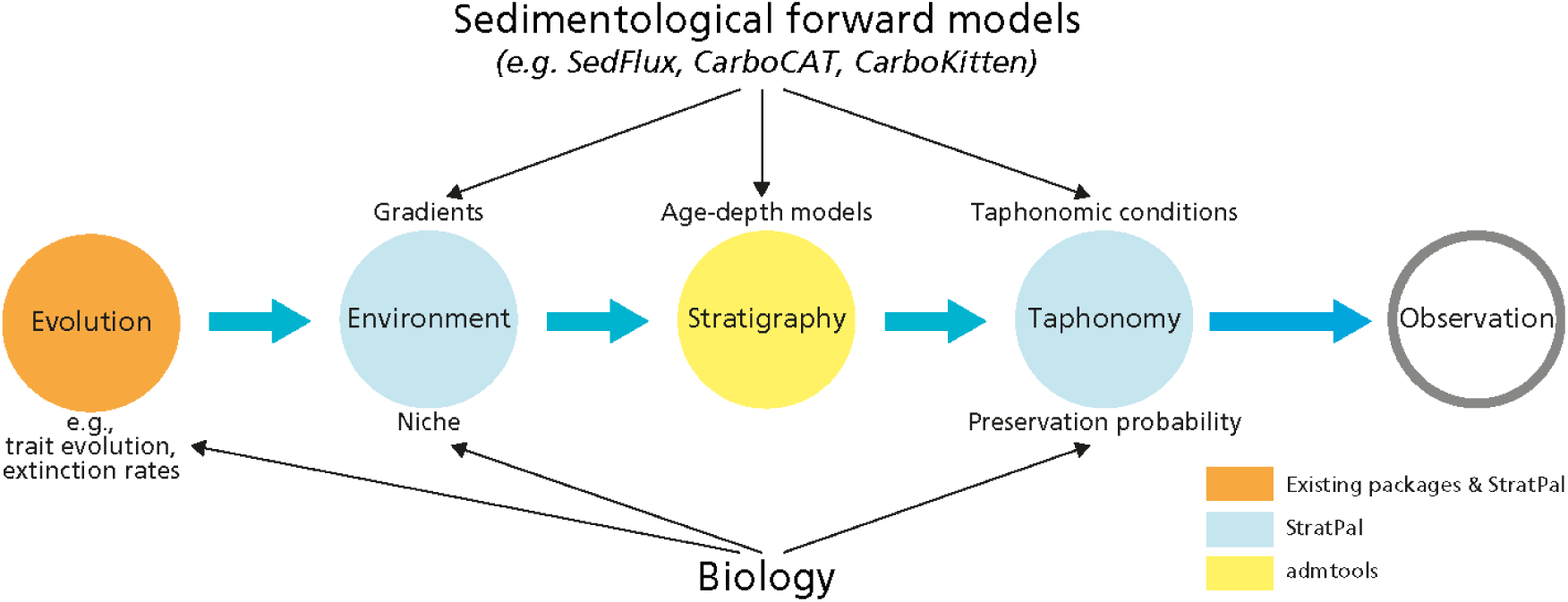
Conceptual framework of the modeling pipeline components. The `StratPal` package models evolutionary change, niches, and taphonomy, the `admtool`s package models stratigraphic biases. Information from sedimentological forward models provide input on gradients, age-depth models, and taphonomic conditions, while biological assumptions are used for the models of evolutionary change, niche definitions, and preservation probability. Note that the left side of the figure is in the time domain, while the right side is in the stratigraphic domain.

### Simulating biological change

Out of the box, `StratPal` comes with options to simulate phenotypic evolution in a lineage and discrete biological events such as first and last occurrences. Details on how user-defined types of biological variables can be implemented are provided in the package vignettes.

For phenotypic evolution, available modes of evolution that can be simulated are stasis (Hunt 2006), strict stasis (Hunt, Hopkins, and Lidgard 2015), (un)biased random walk (Bookstein 1987), and Ornstein-Uhlenbeck processes (Lande 1976) (reflecting convergence to a phenotypic optimum). These modes can be simulated on mean trait level or for individual specimens, providing a connection with the `paleoTS` package (Hunt 2006; 2022).

We refer to “event type data” as any data that consists of discrete events such as occurrences in time or location of individuals or first and last occurrences of taxa. Albeit these events have very different biological interpretations, we treat them together, as they are structurally identical. `StratPal` provides the option to simulate event type data via a constant rate or variable rate Poisson point process (functions `p3` and `p3_var_rate`, respectively), which can be conditioned to return a prescribed number of events. The simulation is based on the procedure described in Streit (2010) p. 14, where the independent and identically distributed (I.I.D.) random numbers are determined using rejection sampling. Poisson point processes assume that events are (mathematically) rare and independent of each other, and the rate prescribes the average number of events per time or length unit.

### Stratigraphic distortions

Stratigraphic distortions are incorporated using the `admtools` package (Hohmann 2024a). This package defines age-depth model objects that serve as coordinate transformation between the time domain (where biological change happens) and the stratigraphic domain (where data is observed). In addition to that, they also contain information on gaps (hiatuses) in the record, allowing to remove data that coincides with times of erosion or non-deposition. The coordinate transformation defined by age-depth models it very general, and can be applied to any process in the time (resp. stratigraphic) domain. This is because it simply replaces the time coordinates with stratigraphic heights (or the other way around) to determine the stratigraphic expression (resp. the temporal expression) of the process. As such, the transformation works equally on time series, event type data representing fossil specimens, first and last occurrences, or phylogenetic trees (time trees). As of now, transformations of phylogenetic trees in the `ape` format (Paradis and Schliep 2019), of lists with temporal or stratigraphic information, of trait evolution series in `pre_paleoTS` format, and of event type data is implemented. Details on how other data types can be transformed can be found in the vignettes of the `admtools` package.

### Preservation

Preservation is modeled using the function `apply_taphonomy`. It combines (1) the change of taphonomic conditions through time or height with (2) the preservation probability of a specimen in dependence of taphonomic conditions to determine how preservation potential changes with time or height. When applied to event type data or lineages, it removes events or specimens according to the preservation probability. Taphonomic conditions can depend on any parameters tracked by the stratigraphic forward model used. Potential examples include, but are not limited to, wave energy, lithology, substrate consistency, or sedimentation rate (Hannisdal 2006).

Note that preservation can be modeled both in the time and the stratigraphic domain depending the parameters determining taphonomic conditions. For lithology, taphonomy would me modeled in the stratigraphic domain, while for wave energy, it would be modeled in the time domain (unless wave energy is itself reconstructed from the rock record). Because of the large number of potential factors and external parameters contributing to taphonomic conditions, we do not provide template functions to construct taphonomic models, but rather provide instructions to implement these models from scratch. This is done by defining two functions representing preservation probability and the change in taphonomic conditions and passing them to `apply_taphonomy`.

### Niche modeling

Niche models can be incorporated using the function `apply_niche`. It combines the niche definition of a taxon with the information about the availability of that niche at a given point in time to determine when a taxon is within its niche and when not, and modifies its abundance accordingly. In the simplest case, a taxon will only occur within the fixed boundaries of its niche (implemented via `bounded_niche`). All specimens/events outside of the boundary will be removed. In a more general case, taxa will be most abundant at specific gradient values, and become less abundant as one moves away from their preferred environmental conditions. To model this, we allow niche definitions where taxa can still exist outside of their preferred environmental conditions, but at a lower abundance. To model this, we represent niches as functions attaining values between 0 and 1, where 0 reflects that the taxon is fully outside of its niche, 1 represents it is fully within its niche, and values in between 0 and 1 indicate the taxon is in its niche, but at reduced abundance. As a template function for this we provide the `snd_niche` function (“snd” standing for “scaled normal distribution”), reflecting the “probability of collection” model from Holland and Patzkowsky (1999). This model uses a bell curve to model niches of marine taxa along a continuous gradient, most commonly water depth.

Note that although conceptually different, modeling niches and taphonomic conditions is mathematically identical: Both are based on the removal of specimens/events depending on a taxon-specific property (niche definition and preservation probability, respectively) and external drivers (change of the ecological gradient or taphonomic conditions with time, respectively).

### Documentation and Open Educational Resources

In addition to the default documentation of functions, the package provides multiple vignettes (long form documentation with extensive worked examples) covering an introduction, stratigraphic paleobiology of event type data and phenotypic evolution, integration with the `paleoTS` package, advanced usage and writing extensions, and documentation of defined `S3` classes. All help pages and vignettes are rendered as a webpage, available under https://mindthegap-erc.github.io/StratPal/. After installation, we recommend going through the available vignettes that explain the functionality of the package in detail.

We provide open educational resources (OERs) for a workshop on building modeling pipelines for stratigraphic paleobiology in R Software. Workshop materials are available via https://doi.org/10.5281/zenodo.13769443 under a CC-BY license and will be continuously updated (Hohmann, Liu, and Jarochowska 2024).

### Development, contribution, and writing extensions

Stable releases of `StratPal` (currently version 0.3.0, Hohmann (2024b)) can be installed from CRAN, development versions are available on GitHub under https://github.com/MindTheGap-ERC/StratPal. The `StratPal` package is an open source project, and we invite other researchers to contribute, contribution guidelines can be found in the CONTRIBUTION.md file in the root of the package GitHub repository. The “Advanced usage and writing extensions” vignette provides background on how this can be done, the “Documentation of defined S3 classes” vignette provides developer documentation. Potential extensions include improving the integration of the package with the existing R package ecosystem for paleontology (e.g., the `FossilSim` (Warnock et al. 2024; Barido-Sottani et al. 2019), `paleotree` (Bapst 2012) or `evoTS` (Voje 2023, 2024) packages), adding vignettes or expanding documentation, or implementing more modes of evolution.

### Application example

All code used for this worked example is available in Hohmann (2024c).

### Example data

The `StratPal` package can be used with outputs from any type of sedimentological forward models, however these models can be challenging to run. We provide example data of stratigraphic architectures with the package to provide meaningful examples and facilitate package usage for users without access to sedimentological forward models. The example data provided is from of a tropical carbonate platform simulated with CarboCAT (Burgess 2013, 2023) under a sinusoidal sea level curve with third and fourth order sea level changes with periods of 1 Myr and 0.112 Myr and amplitudes of 20 and 2 m, respectively. We provide the sea level curve as well as age-depth models, water depths, and stratigraphic columns (bed thickness and facies codes) at 2, 4, 6, 8, 10 and 12 km from shore. This data is extracted from scenario A in Hohmann, Koelewijn, et al. (2024), see therein for simulation details, as well as a chronostratigraphic (Wheeler) diagram and transect of the simulated platform.

As a working example, we examine the formation of fossil accumulations and clusters of last occurrences at different stratigraphic positions, and how this effect varies along the onshore-offshore gradient. This classic example of stratigraphic paleobiology has been simulated in siliclastic systems (e.g., Holland (1995)) and observed empirically (Nawrot et al. 2018), here we show the effect in carbonate systems. More worked examples that show a range of possible applications can be found in the package vignettes, available via `browseVignettes(“StratPal”)` after package installation.

We use two age depth models from the example data set provided with the package, located 2 and 12 km from shore (Figure 2). The age-depth model 2 km from shore is from the platform interior, with more than 145 m sediment accumulated over 2 Myr years and a completeness of 33%. It has 10 hiatuses, of which two are longer that 0.5 Myr and are associated with prolonged drops in sea level. The age-depth model at 12 km from shore is from the proximal slope with 16.5 m accumulated sediment and a completeness of 76%. It has more, but shorter hiatuses (more than 50 hiatuses, median duration 0.048 Myr, 1^st^ and 3^rd^ quartile 0.038 and 0.059 Myr, respectively). This age-depth models is characterized by condensed intervals between 0 and 0.6 Myr (bottom of the section) and 1.4 and 1.8 Myr (14 m height) as indicated by the almost horizontal slope (low sedimentation rate) of the age-depth model.

**Figure 2:**
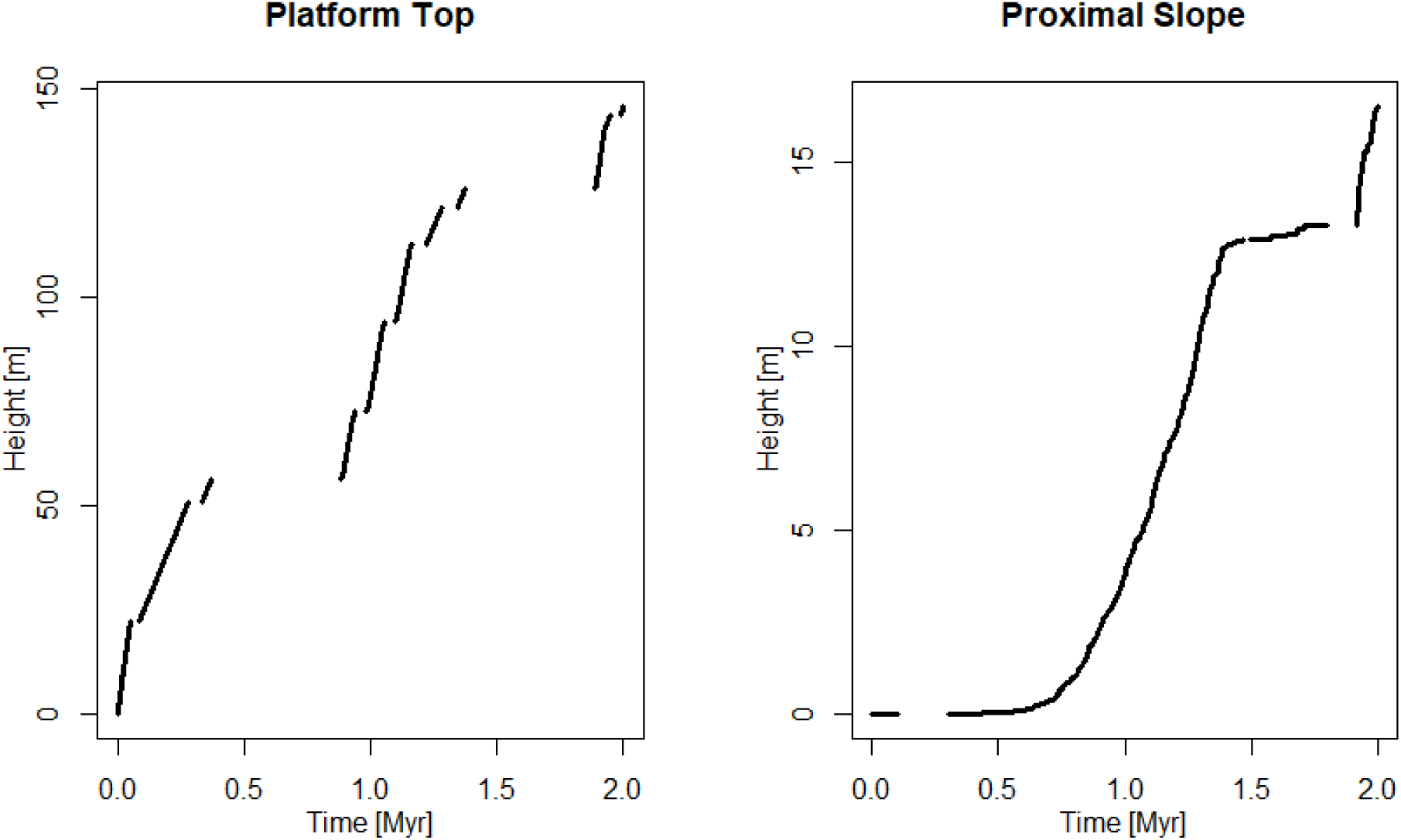
Age-depth models 2 km from shore (platform top) and 12 km from shore (proximal slope) in the simulated carbonate platform provided as example data with the package.

We assume a constant rate of last occurrences and fossil occurrences in the time domain, and examine where stratigraphic distortions lead to cluster formation. We simulate both last occurrences of multiple taxa and occurrences of one taxon as a Poisson point process with a rate of 200 events (last occurrences and fossils, respectively) per Myr via the `p3` function. This high rate was chosen to reduce random fluctuations in the results due to the presence of absence of rare events that can dominate the results under low rates. For last occurrences, we assume that the simulated taxa are sufficiently abundant for the Signor-Lipps effect to be negligible (Signor et al. 1982), meaning the stratigraphic range offset (sampling bias sensu Holland (1995)) is small. See the vignettes on how this assumption can be weakened.

In the platform interior, we observe two clusters of last occurrences coinciding with the stratigraphic positions of the long hiatuses at 56 and 126 m height (Figure 3). This is because under the assumption of abundant taxa, all last occurrences in the time domain that coincide with a gap will be found at the hiatus surface in the stratigraphic domain (unconformity bias *sensu* (Holland 1995)). The lower the abundance of the taxa, the further the cluster will drop below the gap. At the proximal slope, there are to two clusters at 0 and 14 m height (Figure 3). These clusters are associated with stratigraphic condensation, leading to more time (and, as a result, more last occurrences) being represented per stratigraphic interval (Hohmann (2021), condensation bias *sensu* Holland (1995)). When hiatuses and condensation are not recognized, these clusters might be mistaken as elevated extinction rates. Because they are associated with drops in sea level, this might be mistaken for a causal relationship between drops in sea level and elevated extinction rate (“common-cause hypothesis” (Peters and Heim 2011), see also Holland and Patzkowsky (2015)). The clusters in the proximal slope shows that major stratigraphic distortions can also exist in relatively complete sections with short hiatuses given that sedimentation rates vary sufficiently.

**Figure 3:**
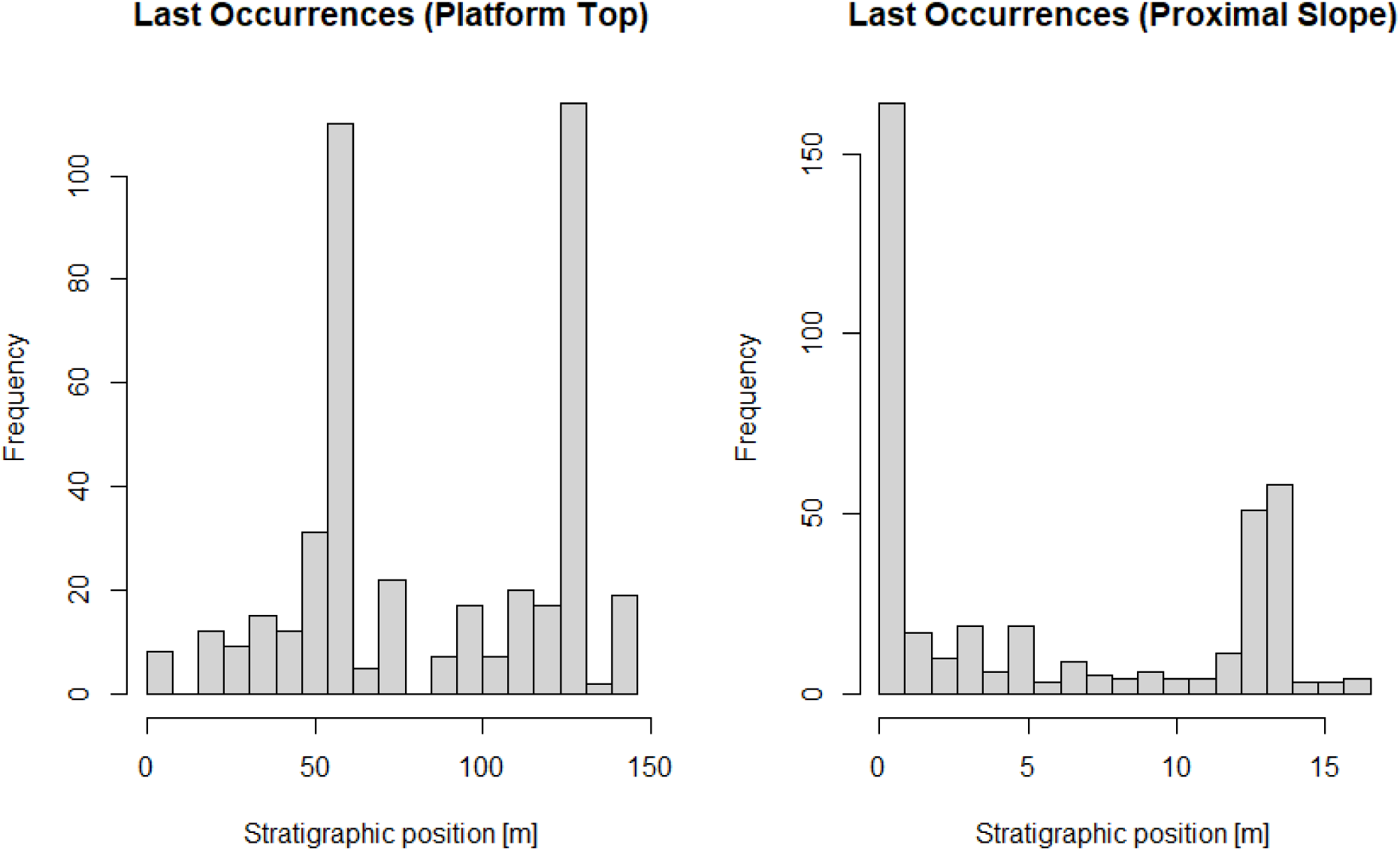
Last occurrences on the platform top (2 km from shore, left) and on the proximal slope (12 km from shore, right) under constant rate of last occurrences in the time domain.

When simulating fossil abundance, we find no significant peaks in fossil abundance on the platform top (Figure 4, left), as fossils coinciding with erosional intervals are destroyed and thus do not cluster at hiatus surfaces (contrary to last occurrences). In contrast, at the proximal slope, we observe fossil accumulations associated with intervals of stratigraphic condensation (Figure 4, right). The contrast between the patterns of last occurrences and fossil abundance on the platform top demonstrate that stratigraphic distortions do not act uniformly, but interact with the primary biological signal.

**Figure 4:**
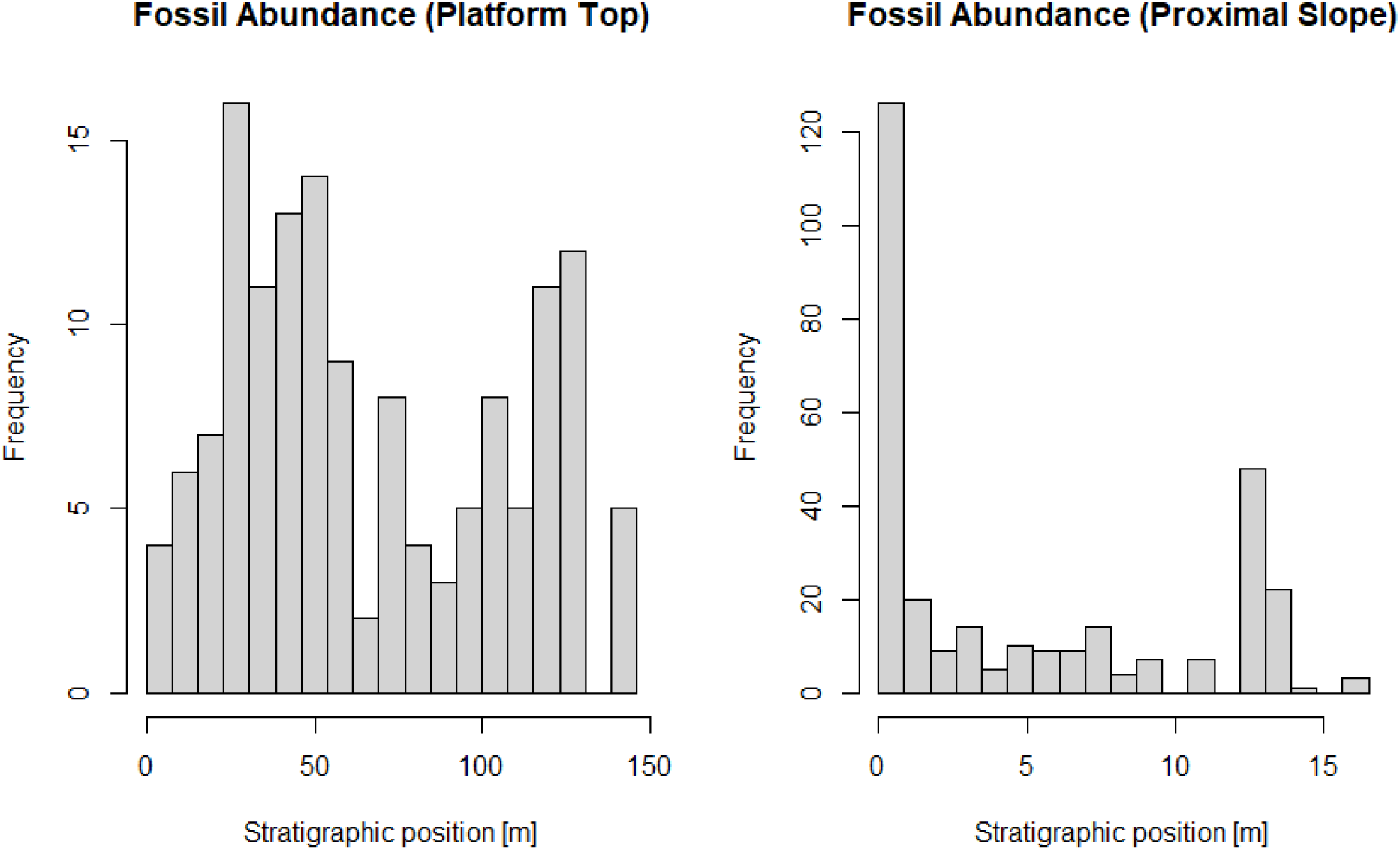
Fossil abundance on the platform top (left) and the proximal slope (right) under constant fossil abundance in the time domain.

In the first example, we implicitly assumed that taxa are equally abundant regardless of environmental conditions. This can be relaxed by using the `apply_niche` function to model the taxon’s preferred water depth, water depth being a good proxy for a number of influential environmental factors for marine organisms. As an example, we use the probability of collection concept of Holland and Patzkowsky (1999) to model the abundance of a taxon that has its optimum water depth at 100 m and is tolerant (eurytopic) with respect to water depth fluctuations.

This reduces fossil abundance towards the top of the section, removing the second peak in fossil abundance. Looking at the water depth at the proximal slope, we can see that this is not a global signal, but rather local extirpation due to the fact that the water depth on the proximal slope increases with time (Figure 5).

**Figure 5:**
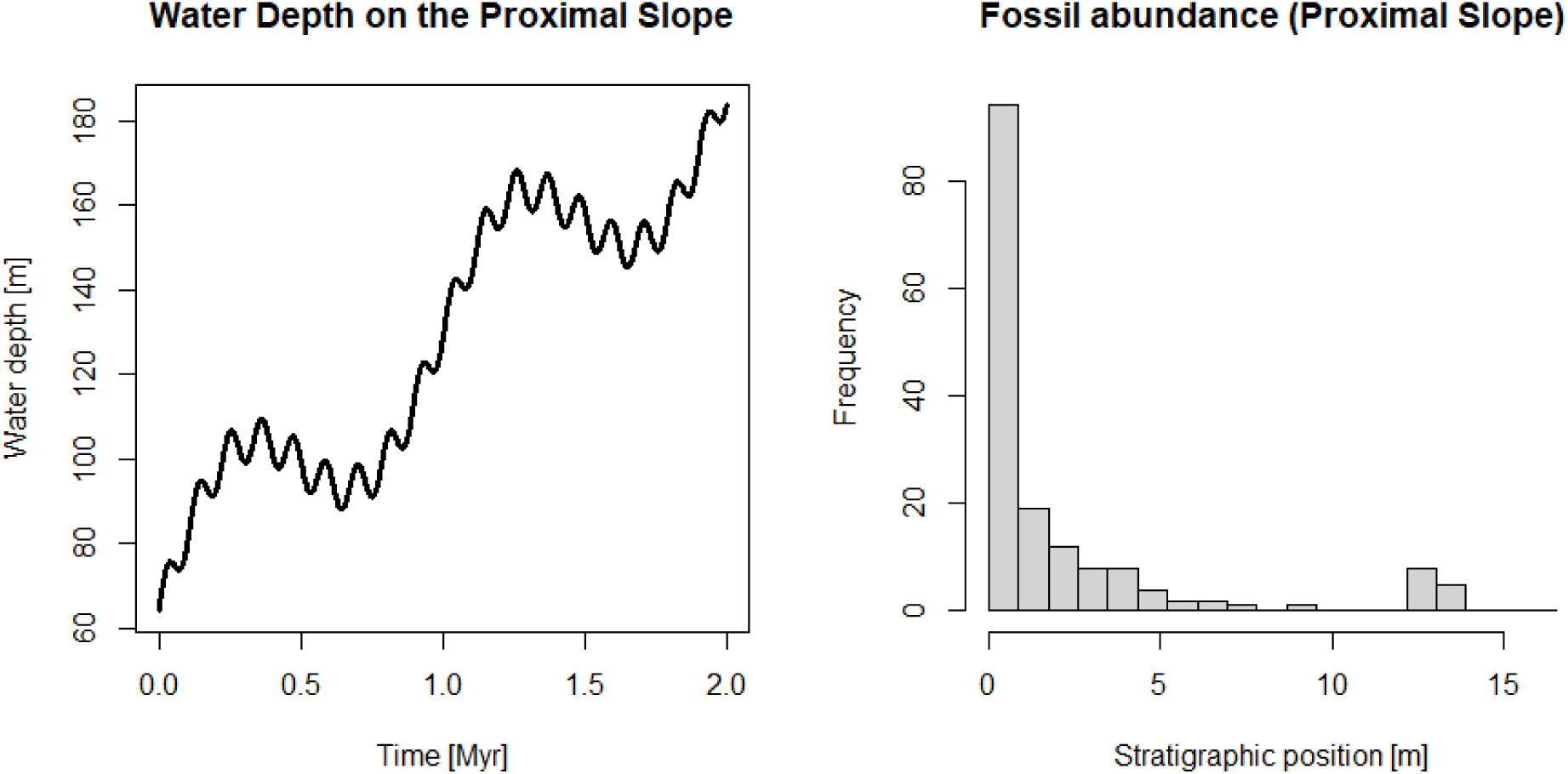
Water depth with elapsed model time and fossil abundance on the proximal slope.

### Application to empirical data

The simple example presented here is derived from simulated data to illustrate the predictive power of the package. However, the toolbox of stratigraphic paleobiology, implemented in `StratPal`, offers the opportunity to evaluate empirical data in diverse ways, including (but not limited to) testing hypotheses on:

1. the stratigraphic expression of fossil occurrences, e.g. by formulating the null hypothesis on the expression in the absence of elevated extinction rate or hypotheses on the expression under different extinction dynamics (Zimmt et al. 2021);
2. taphofacies, i.e. differential preservation of fossils or fossil traits depending on e.g. the lithology or sequence stratigraphic position (Brett 1995; De Baets et al. 2022; Scarponi et al. 2013) by building on prior attempts (Wagner and Marcot 2013) but allowing for inclusion of “taphofacies” i.e. classes of conditions representing different types of preservation, derived from empirical datsets (Holland 2016);
3. the age of fossil occurrences used for calibration of molecular clocks or in phylogenetic inference (Warnock, Yang, and Donoghue 2011; Holland 2016; Barido-Sottani et al. 2019).

Such applications require paleobiologists to adapt more systematically tools already present in the field, but not used systematically: age-depth models, even if very simple, when rates are estimated from empirical data; and the environmental niche. An argument often proposed against the use of age-depth model(s) is that their uncertainty is too big and not enough data is available to formulate one: we show that considering a few end-points (e.g. 100% completeness versus major hiatus) is sufficient to evaluate the robustness of results concerning the rate of extinction and other processes (Hohmann 2021; Hohmann, Jarochowska, and Burgess 2023). As for environmental niches of fossils and their stratigraphic expression, qualitative models have been proposed long before the advent of computer niche modeling, commonly under the term of ecostratigraphy or biofacies (Boucot 1984; Olóriz et al. 1993; Sandberg and Dreesen 1984). These painstakingly compiled models derived from scores of observations yield themselves to formats handled by `StratPal`. For example, the bathymetric niche can be estimated from the frequency of observations reported in the literature and databases (Jarochowska et al. 2017; Parker, McHorse, and Pierce 2018) or from a dedicated empirical investigation (Holland and Zaffos 2011; Patzkowsky and Holland 1999). Overall, the existence of software tools to execute typical analyses in stratigraphic paleobiology should facilitate data collection and study design to answer questions on the preservation of biological patterns in the rock record, without re-implementing all analyses *ad hoc*.

## Conclusion

The `StratPal` package provides a modular, easy to use approach to modeling stratigraphic paleobiology, allowing to build complex models from simple building blocks. Because of its standalone nature, it provides modeling framework where stratigraphic forward models can be used interchangeably, thus opening the possibility to compare the effects of stratigraphic paleobiology across different environments (marine *vs.* terrestrial, siliciclastic *vs.* carbonate *vs.* mixed systems). Each of these systems responds differently to external drivers, resulting in environment-specific preservation of biological signals. The `StratPal` package provides a unified, environment-agnostic approach to model stratigraphic paleobiology, and connects with existing R packages for analytical paleontology. It is openly developed on GitHub under https://github.com/MindTheGap-ERC/StratPal and we invite to contributors to add functionality and improve the integration with existing R package ecosystem for paleontology and paleobiology.

## Code and data availability

All data and code used in the example is available under https://doi.org/10.5281/zenodo.14204077 (Hohmann 2024c).

## Author contributions

Based on the CRediT taxonomy.

Niklas Hohmann: Conceptualization, Formal analysis, Investigation, Methodology, Software, Validation, Visualization, Writing - original draft, review & editing.

Emilia Jarochowska: Funding acquisiton, Project administration, Validation, Writing - original draft, review & editing.

